# Periplasmic proteostasis enables bacterial survival during MreB cytoskeletal disruption

**DOI:** 10.64898/2026.01.06.698069

**Authors:** T. Nagarajan, Muruganandam Nandhakumar, Balaji Kannan, Sharayu Magar, Shanmugapriya Kannaiah, Orna Amster-Choder, Sutharsan Govindarajan

**Author notes:** These authors contributed equally (T. Nagarajan, Muruganandam Nandhakumar). Corresponding author –.

## Abstract

The bacterial actin homolog MreB is essential for various cellular processes, including cell wall biosynthesis, membrane organization, and cell polarity determination. Given its multifaceted roles, MreB is considered a potential target for antibiotic development. However, the bacterial response to MreB inhibition and the factors contributing to bacterial survival under such conditions are not well understood. In this study, RNA sequencing (RNA-seq) was used to identify genes that are differentially expressed in response to MreB inhibition by the A22 antibiotic or to deletion of the *mreBCD* operon. We identified 6 upregulated genes and 24 downregulated genes under both conditions. To determine whether the upregulated genes contribute to bacterial survival during MreB inhibition, we performed A22 antibiotic susceptibility assay on mutants deleted for each of the 6 upregulated genes. Our findings reveal that cells lacking DegP, a periplasmic serine protease, are highly suceptible to A22 treatment. Complementation analysis showed that wild-type DegP, but not a protease-defective mutant, mitigated the effects of A22. The morphological defects in DegP-deficient cells caused by A22 were reduced by ectopic expression of related periplasmic proteases, such as DegQ and DegS. Furthermore, elevated temperatures could alleviate the effects of A22 in a DegP-dependent manner. Overall, our study provides a comprehensive analysis of the global transcriptome-wide effects of MreB inhibition, offering new insights into bacterial response to cytoskeleton disruption. The findings highlight the critical role of DegP in bacterial survival during MreB inhibition and suggest potential avenues for developing novel combinatorial antibiotic strategies targeting MreB and DegP.

**Highlights:**

- MreB disruption by A22 or mutation causes major transcriptome alterations
- Six genes are consistently upregulated under both MreB disruption conditions
- DegP, a periplasmic protease, is crucial for bacterial tolerance to A22
- Elevated temperatures alleviate A22 toxicity in a DegP-dependent manner

## INTRODUCTION

MreB is a bacterial cytoskeletal protein similar to eukaryotic actin, which is highly conserved in many rod-shaped bacteria, including pathogens such as *Escherichia coli*, *Klebsiella pneumoniae*, *Acinetobacter baumannii*, and *Pseudomonas aeruginosa*. MreB controls cell shape, specifically cell elongation, by interacting with cell wall biosynthetic enzymes and regulating the location of cell wall synthesis along the long axis of the cell. Disruption of MreB affects cell wall integrity and cell shape, ultimately resulting in cell death (1,2). MreB forms filaments by polymerization of monomers in an ATP-dependent manner. These filaments are localized underneath the cytoplasmic membrane and form an essential membrane-bound complex together with MreC, MreD, and RodZ (1,3,4). Current evidence suggests that the filaments formed by MreB are short and dynamic, rotating around the long axis of the cell. The movement of MreB filaments is driven by the process of cell wall synthesis. Thus, MreB-cytoskeletal and its associated proteins including MreC, MreD and RodZ are intricately linked for coordinated cell wall synthesis and cell shape maintenance in bacteria (5–7). Besides its well-established role in cell wall synthesis, MreB is also involved in cell division, membrane organization, chromosome segregation, and intracellular transport of molecules (8–10).

The critical role of MreB in bacterial cell shape determination and its involvement in multiple pathways make it an ideal candidate for antibiotic development. Several small molecules targeting MreB have been discovered. Among them, A22, a derivative of S-benzylisothiourea, is a well-known first-generation MreB inhibitor (11,12). Cells treated with A22 or its analog MP265 become spherical due to MreB inhibition (13,14). Biochemical and structural studies have shown that A22 binds near the nucleotide-binding pocket of MreB, affecting ATP-induced conformational changes and resulting in the depolymerization of MreB filaments (15). Second-generation MreB inhibitors like CBR-4830 and third-generation inhibitors like TXH11106, with improved depolymerization activity and a very low frequency of resistance, have been developed as potent antibacterial agents (16,17). These molecules have promising antibacterial effects against important pathogens (17). However, none of these molecules have reached clinical practice due to limitations, including cytotoxic effects, the development of resistance mutations, and potential off-target effects on eukaryotic actin (18,19).

MreB is an essential protein, and its inactivation under normal conditions results in cell death. However, under certain growth conditions or genetic backgrounds, disruption of MreB is tolerated by bacterial cells, as evidenced by their ability to grow in the presence of antibiotics like A22. These conditions include slow growth, high cell density, increased concentrations of Mg2+, the presence of suppressor mutations that overexpress cell division genes such as *ftsZAQ*, and an altered metabolic state resulting in increased gluconeogenesis (20–23). In contrast, the absence of certain other genes results in increased cell death and hypersensitivity to the antibiotic A22 upon MreB disruption. These genes include *envC*, an activator of cell wall amidases (24); *cpxR*, an envelope stress response regulator (25); and *secA* and *secB*, components of the Sec protein translocation pathway (24,26). Thus, a complex interplay of genes, growth conditions, and metabolic status contributes to cellular survival when the essential cytoskeletal protein MreB is disrupted. A deeper understanding of the global response to bacterial MreB inhibition could lead to the development of more effective antibacterial strategies targeting MreB.

In this study, a transcriptomics approach was employed to uncover genes that were differentially expressed during MreB inhibition through two approaches – genetic mutation and antibiotic inhibition. Through global transcriptome analysis and antibiotic survival assays, we found that the periplasmic protease DegP is crucial for bacterial survival during MreB inhibition. These findings reveal a previously unrecognized link between the bacterial cytoskeleton, periplasmic proteases, and the heat shock response, highlighting opportunities to develop future combination strategies to combat pathogens encoding MreB.

## RESULTS

### MreB disruption induces global transcriptome changes

To understand the global response due to MreB inhibition, we performed transcriptome analysis of two different types of MreB-disrupted cells: wild-type cells treated with the MreB-targeting antibiotic A22, and *ΔmreBCD* mutant, which is deleted for the *mreBCD* operon and survives due to suppressor mutations (23). The gene expression profiles obtained in both cases were compared with that of untreated wild-type cells. Before proceeding with RNA extraction and sequencing, samples were verified using phase contrast microscopy. Wild-type cells grown in the presence of 2 µg/ml A22 for 3 hours, but not 1 µg/ml A22, appeared completely spherical, indicating MreB disruption. Similarly, *ΔmreBCD* mutant cells appeared round-shaped, while untreated wild-type cells were rod-shaped **(Fig. 1A)**. Overnight incubation with 2 µg/ml A22 has been found toxic to the cells, as evidenced by relative viability assay **(Fig. S1)**. Hence, we restricted the MreB inhibition to 3 hours. Transcriptome analysis was performed after isolating and sequencing RNA from these samples. A summary of the differentially expressed genes with an adjusted p-value ≤ 0.05, log₂ fold change ≥ 1.0, and maximum count > 20 is presented in **Fig. 1B**, and the corresponding volcano plot is shown in **Fig. 1C**. The complete list of transcriptome-wide changes of all genes is summarized in **Table S3**. The list corresponding to categories of upregulated and downregulated genes in A22, as well as those in the *ΔmreBCD* mutant, are presented in **Table S4**.

**Fig 1.**
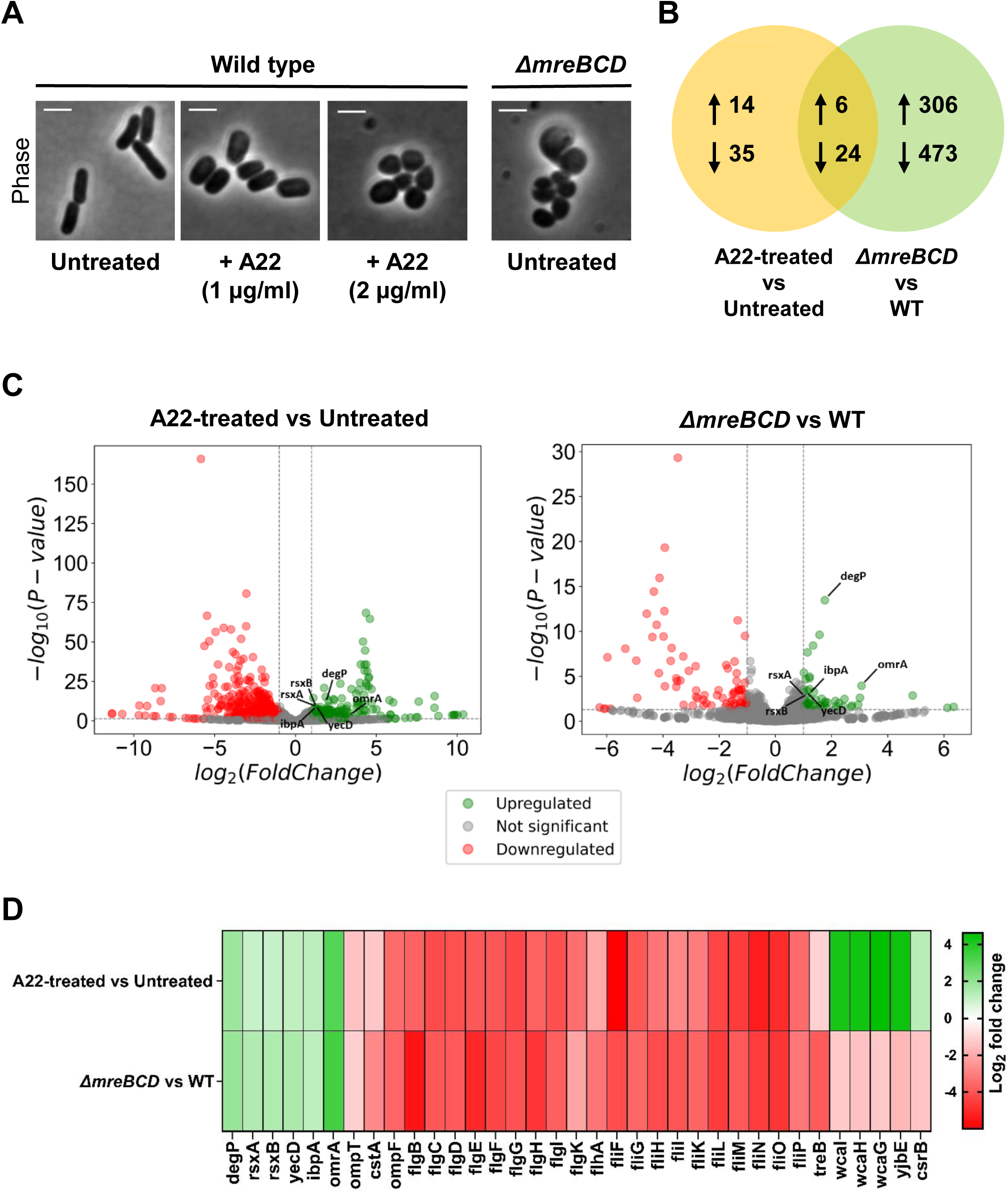
Changes in gene expression upon MreB cytoskeleton inhibition by A22 treatment and *ΔmreBCD* mutation. (A) Phase-contrast images of untreated wild-type (WT), A22-treated WT, and Δ*mreBCD* cells. Scale bar: 2 μm. (B) Venn diagram summarizing the number of differentially expressed genes. (C) Volcano plot illustrating RNA sequencing data. The expression levels of 4,497 genes in both A22-treated vs. untreated and Δ*mreBCD* vs. WT cells were analyzed. The X-axis represents the log₂ fold change values, while the Y-axis denotes the significance level (–log₁₀ p-value). Significantly upregulated genes (adjusted p-value ≤ 0.05) in both A22-treated vs. untreated and Δ*mreBCD* vs. WT conditions are shown in green, while significantly downregulated genes are shown in red. Dotted lines indicate thresholds at p = 0.005 (horizontal) and log₂ fold change = 1 (vertical). (D) Heat map showing the expression patterns of genes differentially expressed in both A22-treated vs. untreated and Δ*mreBCD* vs. WT conditions.

A22-treated cells and the *ΔmreBCD* mutant exhibited distinct transcriptional changes, with some overlaps **(Fig. 1B and Table S4)**. Relative to all *E. coli* genes, only 1.07% were differentially expressed in A22-treated cells, whereas 17.1% were differentially expressed in *ΔmreBCD* mutant cells. A total of 320 genes were upregulated upon MreB disruption, with 14 genes and 306 genes upregulated in A22-treated cells and *ΔmreBCD* mutant cells, respectively. Among these, 6 genes were commonly upregulated in both conditions. Additionally, a total of 508 genes were downregulated upon MreB disruption, with 35 and 473 genes downregulated in A22-treated cells and *ΔmreBCD* mutant cells, respectively. Of these, 24 genes were commonly downregulated in both conditions. The heat map depicts the expression profiles of genes differentially expressed in A22-treated versus untreated and *ΔmreBCD* versus WT cells **(Fig. 1D)**. In total, 35 genes were shared between the two conditions, comprising six commonly upregulated and 24 commonly downregulated genes. Although the majority of these genes displayed similar differential expression trends in both conditions, 5 genes exhibited discordant expression patterns between A22-treatment and *ΔmreBCD* deletion. These data reveal that major transcriptional changes were observed in *ΔmreBCD* mutant cells compared to A22-treated cells. This could be due to the accumulation of suppressor mutations in *ΔmreBCD* mutant cells contributing to their survival.

### Analysis of differentially expressed genes upon MreB disruption

KEGG Pathway enrichment analysis was performed using the DAVID database to further understand the transcriptional response to MreB inhibition. The significantly enriched pathways are summarized in **Fig. 2A**, and the analysis data are provided in **Table S5.** In the *ΔmreBCD* mutant, the exopolysaccharide biosynthesis, aminoacyl-tRNA biosynthesis, and arginine biosynthesis pathways were upregulated. In contrast, in A22-treated cells, only a few genes involved in fewer pathways, particularly motility pathways and the histidine biosynthetic process, were upregulated. Downregulated genes in *ΔmreBCD* cells correspond to several pathways, including oxidative phosphorylation, flagellar assembly, and multiple metabolic pathways, including carbon metabolism. Notably, 18 genes of the tricarboxylic acid cycle (TCA) and 16 genes of the phosphotransferase system (PTS) were downregulated in these cells. In the case of A22-treated cells, 20 genes of flagellar assembly and 3 genes of chemotaxis were downregulated **(Table S5).** Recently, the TCA cycle and the PTS system were implicated in the resistance of bacterial cells towards MreB disruption (21,24). Similarly, MreB disruption was previously shown to downregulate the expression of flagella genes in *Salmonella* (27). The similarity to the changes observed here validate the transcriptomic data.

**Fig 2.**
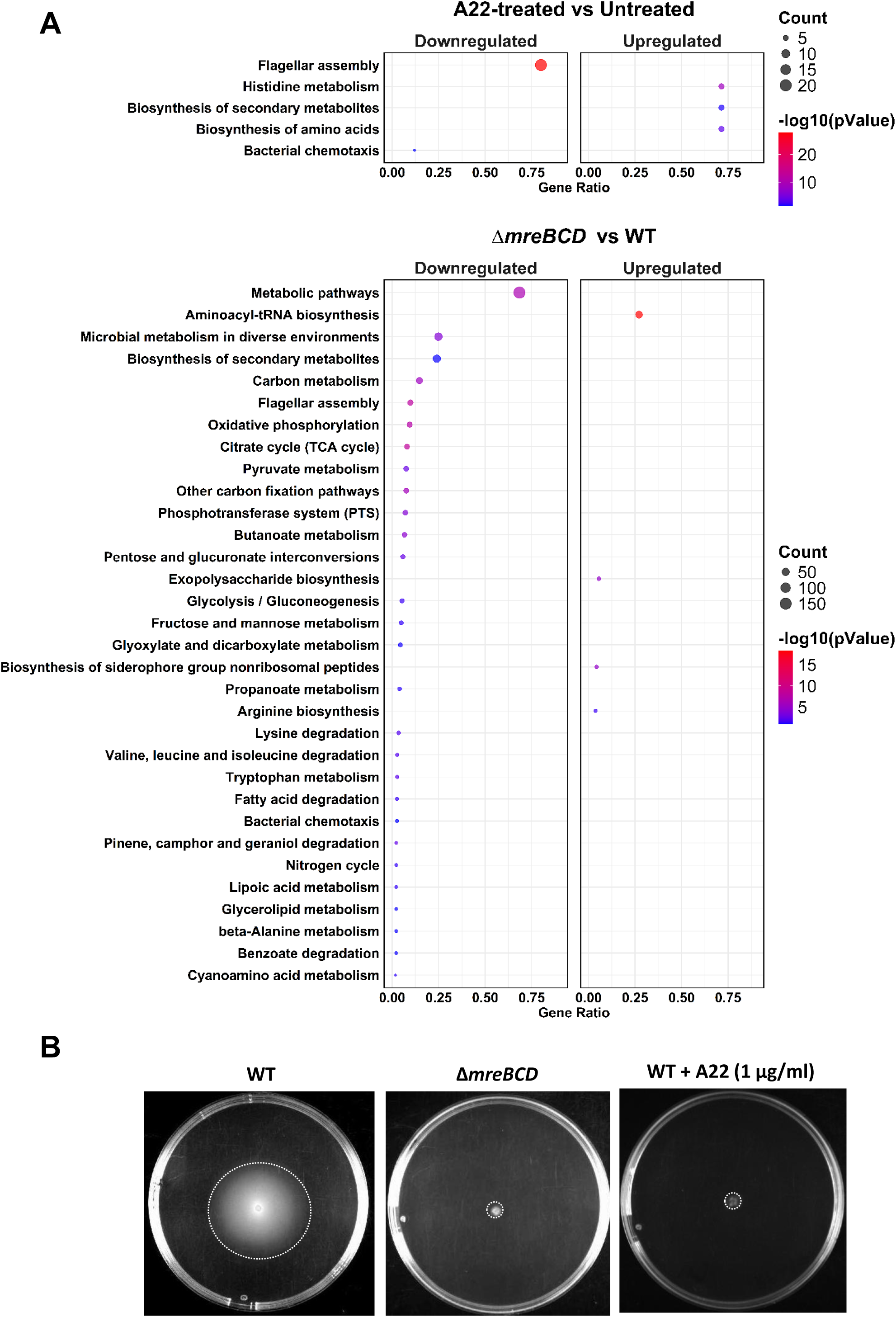
KEGG pathway enrichment analysis and motility assay. (A) Dot plot illustrating the distribution of KEGG pathway gene sets among all differentially expressed genes during MreB inhibition induced by A22 treatment or the Δ*mreBCD* mutation. The color gradient of dots represents the significance of enrichment (-log_10_(p-value)). In contrast, the size of each dot indicates the number of genes from the dataset involved in the pathway (Count). The X-axis represents the gene ratio, a proportion of genes from the dataset that are associated with a particular KEGG pathway. (B) Colonies of WT, Δ*mreBCD* mutant, and WT + A22 (1 µg/ml) were grown for 12 hours on soft agar plates.

To verify whether downregulation of the flagellar assembly and chemotaxis pathways under both MreB disruption conditions are manifested by physiological changes, we tested the motility of wild-type and MreB-disrupted cells. We observed that MreB-disrupted *E. coli* cells, both via A22 treatment and *ΔmreBCD* mutation, are affected in motility **(Fig. 2B)**, consistent with the transcriptomic data. Taken together, these data indicate that the transcriptomic changes observed due to MreB disruption are significant and consistent with observed phenotypic changes.

### Functional analysis reveals DegP’s role in A22 Sensitivity

Based on the transcriptomic data, which provided us with a global overview of changes associated with MreB disruption, we set out to identify specific factors contributing to bacterial survival during MreB disruption. To this end, we focused on the six common factors upregulated under both conditions of MreB disruption **(Fig. 3A).** To understand their functional importance during MreB disruption, we constructed single-gene deletions of each of these genes in the MG1655 background and compared the survival of these mutants with the wild-type in the presence of sub-inhibitory concentrations of A22 (1 µg/ml) through spot-titre analysis. The results, compiled in **Fig. 3B**, show that the deletion of *degP* severely compromised viability in the presence of A22, indicating its crucial role in antibiotic tolerance. In contrast, the other mutants exhibited similar viability to the wild-type, suggesting that their increased expression during MreB disruption is not required for bacterial survival **(Fig. 3B).** Since DegP is part of the cellular protein homeostasis machinery, we wanted to know whether the observed effect is specific to DegP or can be observed in other proteolytic mutants. A22-sensitivity analysis of *Δlon* and *ΔclpX* cells, which are defective for cytoplasmic proteolytic systems, did not show any difference compared to wild-type, indicating that the observed effect is specific to DegP-defective cells **(Fig. 3C).** Consistent with the A22 hypersensitivity observed in the *ΔdegP* mutant, microscopic analysis showed that *ΔdegP* cells became round-shaped upon exposure to sub-inhibitory concentration of 1 µg/ml of A22, whereas wild-type cells were only slightly affected in terms of cell shape **(Fig. 3D).** These results suggest that the DegP-associated periplasmic stress pathway is a critical factor for bacterial survival during MreB inhibition, and we decided to explore this further.

**Fig 3.**
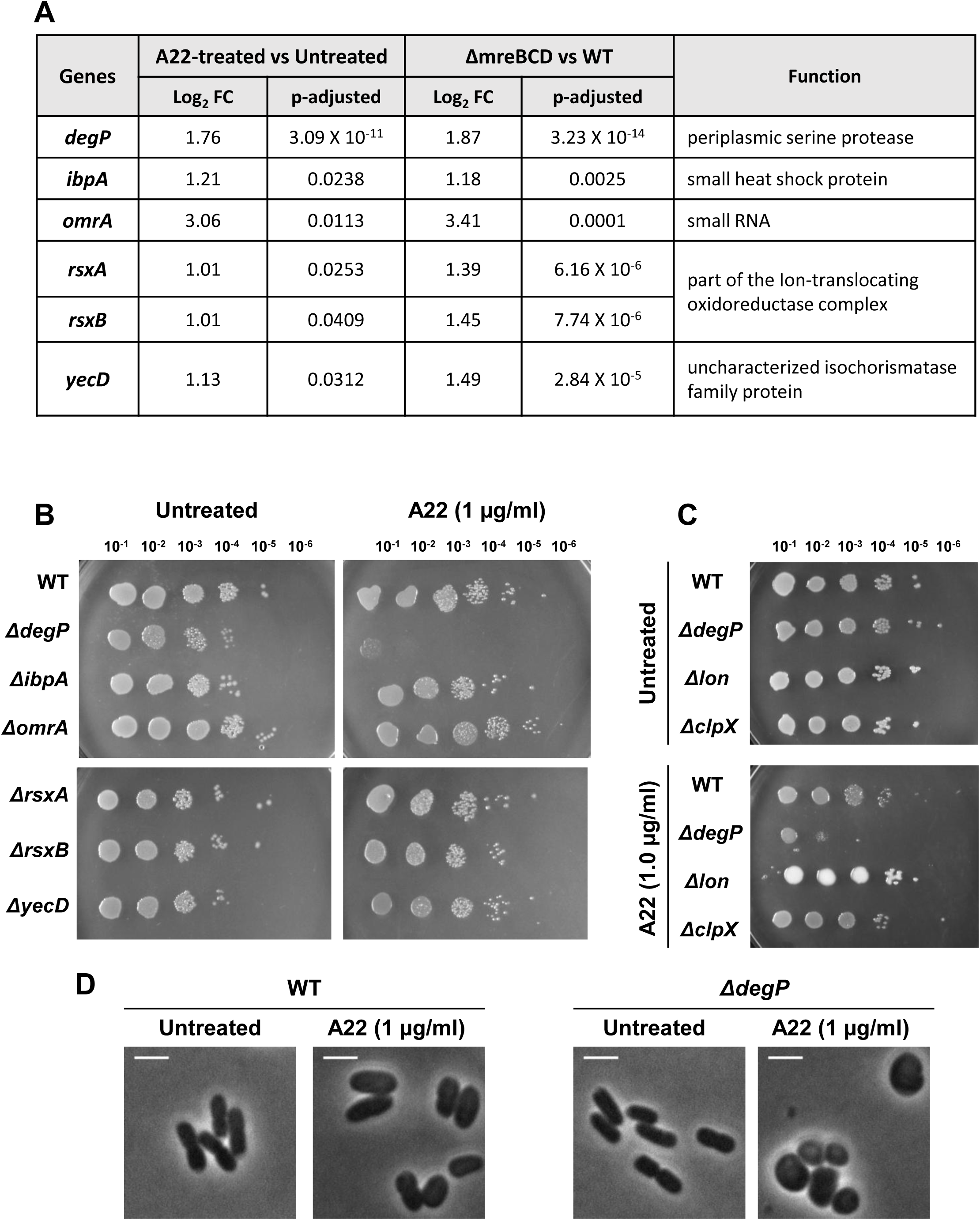
Periplasmic protease DegP is critical for bacterial survival during A22 treatment. (A) Table listing genes that are significantly upregulated by A22 treatment and the Δ*mreBCD* mutation. (B, C) Serial dilutions of the indicated strains were grown on LB plates without or with the antibiotic A22 (1 µg/ml). (D) Phase-contrast images of wild-type and Δ*degP* cells exposed to A22 (1 µg/ml) for 3 hours. Scale bar: 2 μm.

### Contribution of periplasmic proteolytic pathway to survival during MreB disruption

DegP is a major periplasmic serine protease, which removes aberrant proteins from the periplasmic space (28,29). The impaired survival of Δ*degP* mutant cells in the presence of A22 suggests periplasmic protein misfolding issues during MreB disruption. DegP, DegQ and DegS are the three high temperature requirement (HtrA) proteases which play redundant roles in periplasmic proteolytic process (30). To validate the importance of this pathway in A22 antibiotic sensitivity, we cloned wild-type DegP, DegQ, and DegS in a plasmid after an inducible promoter. In addition, a protease defective mutant version of DegP which harbours an amino acid change in position 210 (S210A), was also cloned and designated as DegP*. In complementation analysis, we observed that temperature sensitivity phenotype of Δ*degP* mutant could be rescued by expression of DegP and DegQ, but not DegP* or DegS **(Fig. S2),** consistent with previous reports (30,31).

Next, we assessed the ability of the plasmid-encoded DegP, DegQ, DegS, and DegP* in alleviating the A22 hypersensitivity of Δ*degP* cells. Spot titre assay shows that expression of plasmid-encoded DegP, DegQ, or DegS, in the presence of the inducer arabinose, exhibited a slight 10-fold increase in viability of Δ*degP* cells in the presence of A22 **(Fig. 4A).** In contrast, protease-defective DegP* expressing cells did not show any difference in the viability and appeared similar to the control **(Fig. 4A).** Furthermore, morphological analysis clearly demonstrated the role of Deg proteases in mitigating the effects of A22. Specifically, plasmids expressing DegP, DegQ, and DegS—but not the DegP* mutant—were able to markedly reduce the cell rounding phenotype caused by A22 treatment of Δ*degP* cells **(Fig. 4B)**, suggesting that increased expression of periplasmic proteases can counteract MreB disruption, possibly by degrading the misfolded targets of DegP and thereby reducing the burden of growth defect.

**Fig 4.**
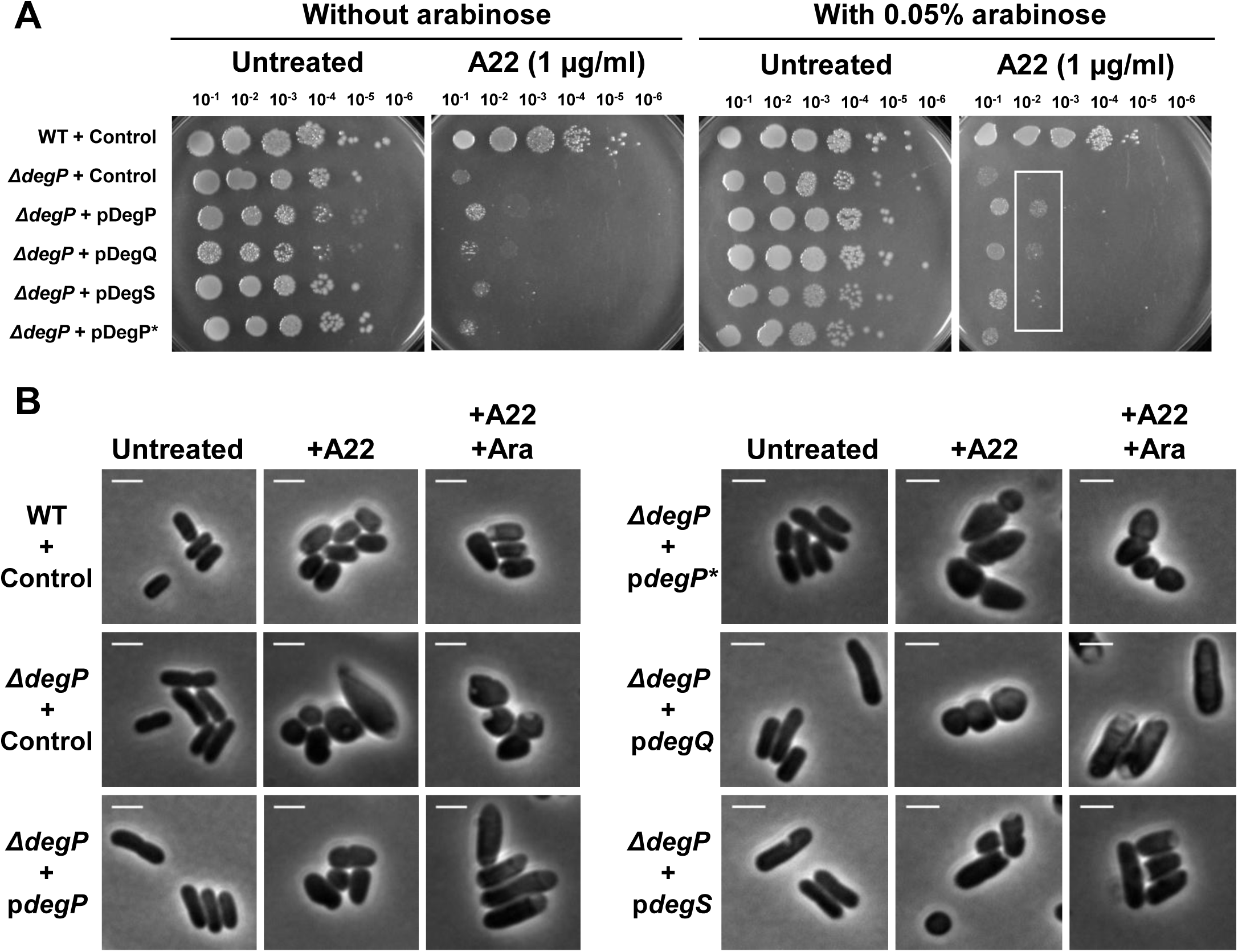
Plasmid complementation of periplasmic proteases moderately alleviates the A22 hypersensitivity of Δ*degP* mutant cells. (A) Serial dilutions of Δ*degP* cells transformed with an empty vector control or plasmids expressing DegP, DegQ, DegS, or DegP*. Serial dilutions were spotted on LB plates without or with the inducer arabinose (0.05%) and incubated overnight at 37°C. One set of plates contained A22 (1 µg/ml), while another was left untreated. Expression of DegP, DegQ, or DegS improved the growth of Δ*degP* cells by approximately tenfold, as indicated by the white box. (B) Phase-contrast images of Δ*degP* cells containing the empty vector control or plasmids expressing DegP, DegQ, DegS, or DegP*, grown in the presence or absence of arabinose (0.05%). One set of strains was treated with A22 (1 µg/ml), and another was left untreated. Scale bar: 2 µm.

### Elevated temperature alleviates survival during MreB inhibition in a DegP-dependent manner

DegP-associated periplasmic proteases are induced as part of the heat shock response to remove misfolded proteins in the periplasm (32). Hence, we wanted to explore the possible reciprocity between the cytoskeleton and heat shock. To this end, we assessed the survival of wild-type cells exposed to a toxic concentration of A22 (2 μg/ml) at three different temperatures: 30°C, 37°C, and 42°C. Strikingly, incubation at higher temperatures significantly mitigated the toxic effects of A22 compared to lower temperatures **(Fig. 5A and 5B).**

**Fig 5.**
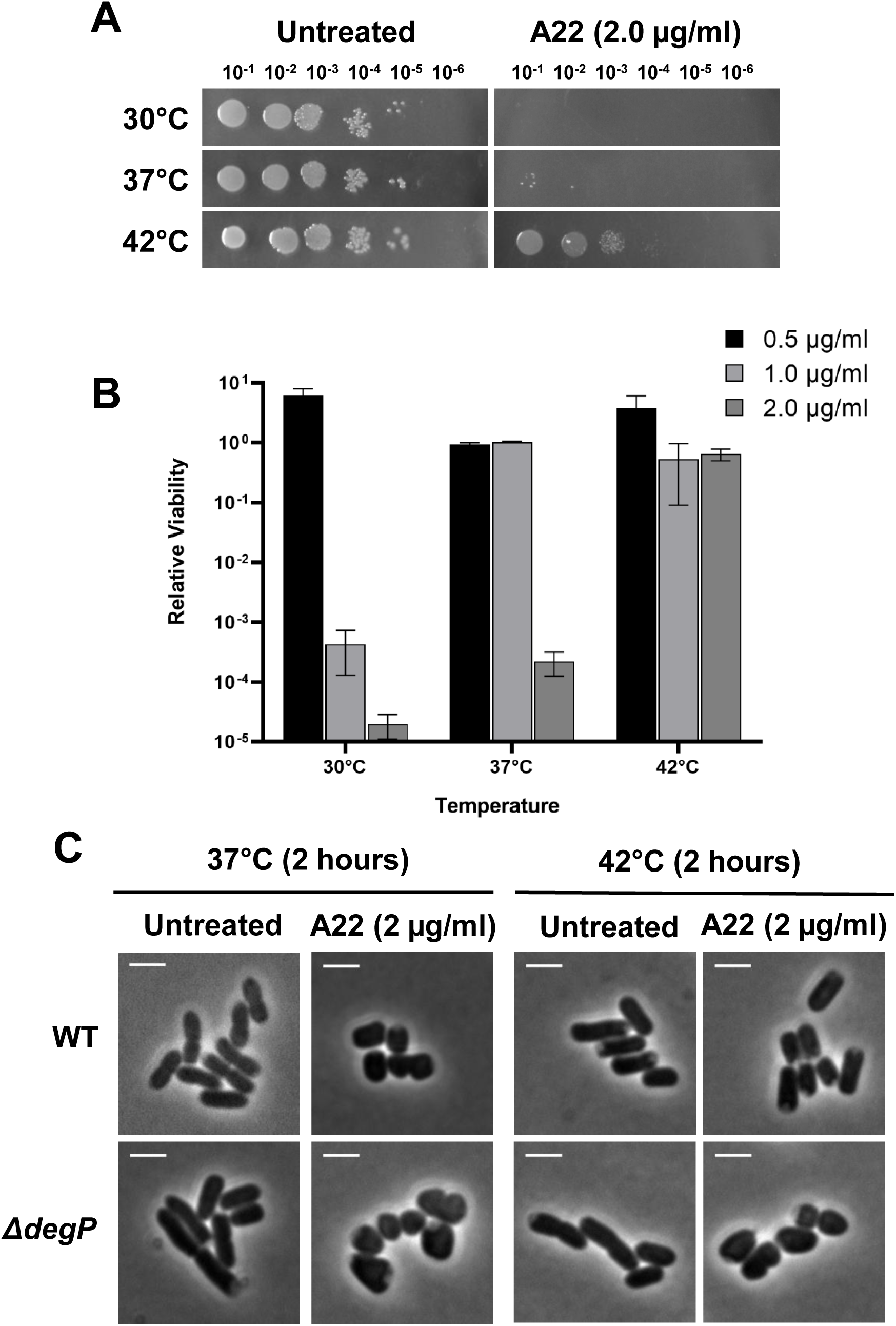
Elevated temperatures alleviate A22 toxicity in a DegP-dependent manner. (A) Serial dilutions of WT cells grown on LB plates with or without the antibiotic A22 (2 µg/ml) and incubated at 30°C, 37°C, or 42°C overnight. (B) Bar graph showing the relative viability of WT cells grown at various concentrations of A22 and incubated at 30°C, 37°C, or 42°C overnight. (C) Phase-contrast images of WT and Δ*degP* cells exposed to A22 (2 µg/ml), either untreated or treated with A22 (2 µg/ml) and grown at 37°C or 42°C for 2 hours. Scale bar: 2 µm.

We next investigated the role of DegP in the alleviation of A22 toxicity at elevated temperatures. As *ΔdegP* cells are temperature sensitive, we could not perform a growth assay to evaluate their survival in the presence of A22 at 42°C. As an alternative, we employed morphological assessments comparing wild-type and *ΔdegP* cells following short-term exposure (2 hours) in the presence of A22 (2 μg/ml) to 37°C and 42°C. The results presented in **Fig. 5C** show that, wild-type cells exposed to A22 at 42°C largely maintained their rod shape as opposed to exposure at 37°C. In contrast, *ΔdegP* cells failed to maintain rod shape in the presence of A22 at both temperatures **(Fig. 5C).** This suggests that elevated temperatures alleviate A22 toxicity in a DegP-dependent manner.

## Discussion

Components of the bacterial cell envelope are common targets of antibiotics, with β-lactams being a well-characterized class that inhibit peptidoglycan synthesis by binding to penicillin-binding proteins (PBPs) (33,34). This inhibition compromises cell wall integrity and leads to cell lysis. The small molecule antibiotic A22, in contrast, disrupts the cell envelope by targeting MreB, the bacterial actin homolog and a central cytoskeletal protein localized to the cytoplasmic side of the inner membrane (11,12,15). MreB polymers exhibit complex association with the cell wall synthesis machinery, and their inhibition leads to uncoordinated cell wall synthesis and eventually cell lysis (22). Beyond impairing cell wall integrity, MreB disruption also results in defects in chromosome segregation, lipid homeostasis, and protein localization (8–10), suggesting that targeting MreB can lead to pleiotropic effects on essential cellular processes.

Our transcriptomic analysis of MreB inhibition provides a glimpse to the global gene expression changes that occur during MreB disruption. Gene expression changes are more pronounced in the *ΔmreBCD* knockout strain compared to wild type cells treated with the antibiotic A22. We propose that the variation in transcriptional response arises from the severity of the *ΔmreBCD* mutation, which is coupled to accumulation of suppressor mutations that enable the survival of this otherwise lethal mutant, since MreB is an essential gene. The suppressor mutations allow slower growth or increased expression of the tubulin homolog FtsZ, allowing survival with poor growth (20,23). Our transcriptomic data points at several differentially expressed genes that can be linked to the survival of the *ΔmreBCD* mutant. For example, we observed significant upregulation of genes involved in cell wall biosynthesis, including *mrdA* (PBP2), *dacD* (PBP6b), and *mrcA* (PBP1a). MrdA (PBP2) is a peptidoglycan DD-transpeptidase, catalyzing the formation of 4-3 cross-links in the peptidoglycan structure, crucial for coordinating cell growth and shape maintenance under stress conditions(35). DacD (PBP6b) is a membrane-bound D-alanyl D-alanine carboxypeptidase that contributes to the maintenance of cell shape at low pH (36). MrcA (PBP1A) is a bifunctional peptidoglycan synthase with transglycosylase and transpeptidase activity and is known to mediate cell wall synthesis independent of MreB (37). Expression of these genes is higher compared to *ftsZ*, which is known to support the survival of this mutant (23). Intriguingly, expression of *mrcB* (PBP1b) was not significantly upregulated, despite prior evidence indicating its importance for bacterial survival during MreB disruption (23). However, since PBP1a is functionally redundant with PBP1b—both possessing transglycosylase and transpeptidase activities (37) —its increased expression in the *ΔmreBCD* mutant likely compensates for the higher expression of PBP1b, thereby potentially supporting cell growth. Given that these genes are key PBPs involved in peptidoglycan crosslinking and synthesis, their elevated expression likely helps maintain cell wall integrity in the absence of MreBCD.

Additionally, downregulation of specific pathways may further contribute to the *ΔmreBCD* mutant’s survival. We observed reduced expression of genes involved in the TCA cycle and sugar transporters (PTS system), both of which are central to energy production and rapid growth. Suppression of these energy-generating pathways likely imparts a slower growth rate, which is known to support the survival of bacteria lacking MreB. Moreover, this repression aligns with recent studies that link metabolic state to resistance to MreB disruption (21,24).

In contrast to the *ΔmreBCD* mutant, A22-treated cells exhibited a more modest transcriptional shift, primarily affecting genes involved in histidine biosynthesis and envelope stress, which may reflect a transient stress response rather than a long-term adaptation. The reason for the upregulation of the histidine biosynthesis pathway during MreB disruption remains unclear. Interestingly, the *hisO1243* mutation, which causes overproduction of HisF and HisH and leads to severe cell division and elongation defects, can be suppressed by mutations in *envB* - a gene later identified as *mreB* (38). This suggests a functional link between histidine biosynthesis and the MreB cytoskeletal system. The observed upregulation of the histidine pathway during MreB disruption may therefore reflect a compensatory or interacting mechanism involving cell shape regulation, envelope stress, connecting with histidine metabolism. However, this connection needs to be studied further. Furthermore, genes related to motility and chemotaxis were markedly downregulated under both conditions. A similar trend has been reported in *Salmonella*, where MreB disruption results in motility defects and downregulation of motility-related genes (27). Another study observed that deletion of a 13 kb genetic region encompassing motility and chemotaxis genes reduces susceptibility to A22 (24). Together, these findings offer a comprehensive view of the pathways differentially regulated in the *ΔmreBCD* mutant and highlight mechanisms that may support its survival and growth in the absence of the MreBCD cytoskeletal system. Still, we observed a limited overlap in differentially expressed genes between the two conditions, potentially due to the chronic nature of the *ΔmreBCD* mutant, which requires several pathways to be differentially regulated for continued survival, in contrast to A22-treated cells, which showed an acute response to short term antibiotic treatment.

DegP is a heat shock-inducible serine protease and chaperone known to function at the interface of protein quality control and stress response in the periplasm, particularly under conditions of envelope stress (39). The hyper susceptibility of Δ*degP* mutants to A22 suggests that MreB inhibition increases the burden of misfolded proteins or cell envelope stress signals that require DegP-mediated clearance. A recent study indicates that A22 treatment causes *E. coli* cells to widen, leading to reduced periplasmic space, which activates the Rcs envelope stress system (40). Misfolded proteins in the periplasm may originate from multiple pathways, most likely involving penicillin-binding proteins (PBPs), whose assembly and proper localization are disrupted during MreB inhibition. Other possible contributors include the misfolding or mislocalization of proteins such as periplasmic MreC, outer membrane proteins, or integral membrane proteins that depend on appropriate lipid environments for correct folding. Thus, the inability of DegP protease-defective cells to cope with the envelope stress response activated during MreB inhibition contributes to the lethal phenotype and underscores the essential role of DegP’s proteolytic activity. Furthermore, the partial rescue observed upon plasmid-mediated expression of DegQ, DegS, or under elevated temperatures suggests functional redundancy and heat shock-induced compensation (30). These findings highlight how cytoskeletal disruption can destabilize envelope integrity, making cells increasingly dependent on DegP to manage stress and maintain cellular homeostasis. Together, our findings suggest that MreB function is coordinated with periplasmic proteostasis, underscoring an integrated cytoskeletal–envelope homeostasis network. Our results open the possibility of co-inhibiting MreB and DegP as a promising strategy to enhance antibacterial efficacy by simultaneously targeting cytoskeletal integrity and stress response mechanisms. Further studies in this direction may lead to the development of novel strategies for combating bacterial infections.

## MATERIALS AND METHODS

### Bacterial strains and growth media

Strains and plasmids used in this study are listed in supplementary Table S1. LB agar and LB broth (containing 1% tryptone; 0.5% yeast extract; 0.5% NaCl) were used for all the experiments. Overnight *E. coli* cultures were grown in LB supplemented with appropriate antibiotics. Overnight cultures of *ΔdegP* strains were grown at 30°C and experiments were conducted at the indicated temperatures. When appropriate, antibiotics for *E. coli* cultures were added at the following concentrations: ampicillin (100 μg/ml), kanamycin (30 μg/ml). All the antibiotics used in this study were purchased from HiMedia (Mumbai, India).

### Construction of plasmids

Plasmids expressing DegP, DegQ and DegS were constructed by Gibson assembly in the pBAD18 plasmid as previously described (41). DegP was amplified using the primers F-RBS-DegP, R-DegP; DegQ was amplified using F-RBS-DegQ, R-DegQ and DegS was amplified using F-RBS-DegS, R-DegS from the genomic DNA of *E. coli* MG1655 K12. The resulting amplicons were PCR purified and assembled using Gibson assembly protocol in the pBAD18 vector backbone, which was amplified using F-pBAD18-gib and R-pBAD18 gib primers.

DegP*, which is a protease defective mutant of DegP harbouring an amino acid change in position 210 (S210A), was constructed as follows: DegP was amplified as two fragments, using four primers, which includes two mutated primers (F-DegP-S210A and R-DegP-S210A), for introduction of the mutant allele. Fragment 1 was amplified using F-RBS-DegP and R-DegP-S210A and fragment 2 was amplified using F-DegP-S210A and R-DegP. Both the fragments were purified and DegP* was constructed using an over lapping PCR using both the fragments as template. The final PCR product is purified and assembled using Gibson assembly protocol protocol in the pBAD18 vector backbone which was amplified using F-pBAD18-gib and R-pBAD18 gib primers. The oligonucleotides used in this study are listed in **Table S2** and were purchased from Eurofins Scientific. All constructs were verified by sequencing.

### RNA Extraction

Total RNA was prepared using the Direct-zol™ RNA Miniprep Kit (Zymo Research) according to the manufacturer’s protocol. Briefly, one volume of ethanol (95–100%) was added directly to one volume of sample homogenate (1:1) in TRI Reagent®, and the mixture was vortexed thoroughly. The resulting mixture was loaded into a Zymo-Spin™ IIC Column placed in a Collection Tube and centrifuged for 1 minute. For samples exceeding 700 µL, this step was repeated. The column was then transferred to a new Collection Tube, and the flow-through was discarded. Next, 400 µL of Direct-zol™ RNA PreWash was added to the column and centrifuged for 1 minute; this step was repeated once. Subsequently, 700 µL of RNA Wash Buffer was added, followed by centrifugation for 1 minute. The flow-through was discarded, and the column was centrifuged for an additional 2 minutes in an emptied Collection Tube to ensure complete removal of the wash buffer. The column was then carefully transferred into an RNase-free tube. For elution, 50 µL of DEPC-treated water was added directly to the column matrix and centrifuged for 1 minute. For highly concentrated RNA, ≥25 µL of elution volume was used. The eluted RNA was either used immediately or stored at ≤−70 °C until further use.

### Library Preparation and Sequencing

Libraries were generated essentially following the RNAtag-Seq approach (42). Briefly, 400 ng of RNA from triplicate samples were fragmented at 94 °C for 3 min, depleted of DNA, dephosphorylated, and purified using RNAClean XP beads. 3′ barcoded adapters were then ligated to the RNA samples, after which barcoded RNAs were pooled, and rRNA was depleted using the Ribo-Zero kit. First-strand cDNA synthesis was performed, followed by degradation of the excess RNA after reverse transcription. Primer dimers were removed using RNAClean XP beads. A second 3′ linker was ligated to the cDNA samples, and unligated adapters were removed using RNAClean XP beads. The resulting libraries were PCR amplified and sequenced on an Illumina HiSeq 5000 platform.

### Analysis of RNA-Seq Data

For RNAtag-Seq data, reads from each sample within a pool were demultiplexed using custom scripts based on their associated barcodes. Then, reads were aligned to the *Escherichia coli* MG1655 genome (RefSeq accession NC_000913) using BWA(43). Read counts were assigned to genes using custom scripts. Differential expression analysis was performed with EdgeR (44) using the likelihood ratio test. Visualization of raw sequencing data and coverage plots in the context of genome sequences and gene annotations was carried out using GenomeView (45). The expression levels of 4,497 genes were analyzed in A22-treated vs. untreated cells and *ΔmreBCD* vs. wild-type (WT) cells, and the results were visualized as volcano plots using Bioinfokit v2.0.8. Genes with an adjusted p-value ≤ 0.05, a log₂ fold change ≥ 1.0, and a maximum count > 20 were considered significantly expressed. This list of genes was subsequently analysed for KEGG pathway enrichment using DAVID Bioinformatics Resources with the default parameters (EASE threshold = 0.1 and count threshold = 2) (46,47). The enrichment results were visualized using the ggplot2 package in R.

### Strain construction through P1_vir_ transduction

P1_vir_ mediated transduction was performed as previously described (48,49). Briefly, the P1vir lysate prepared from the relevant donor strains for each cross **(Table S1)** was used to transfer the relevant allele/marker into recipient strains, and the transduced cells were then plated on appropriate selective plates containing antibiotics. The plates were incubated at 30°C or 37°C until transductants appeared. The obtained transductants were segregated by streaking on LB agar plates containing the appropriate antibiotics. The representative transductants in each case were purified and stored for further analysis. The recipient and donor strains are listed in **Table S1**.

### Relative viability analysis

To quantitatively analyse the survival of bacterial cultures in the presence of A22, a relative viability assay was done as previously described in (48,49). Overnight cultures were sub-cultured into fresh LB broth and grown till mid-log phase. The samples were then diluted in fresh 0.85% saline solution, and appropriate dilutions were spotted on LB plates (to know the cfu/ml) and on LB plates containing A22 (to know the survival). For plasmid-mediated expression, arabinose was added to the LB agar in different concentrations or not added. All the plates were incubated at appropriate temperatures and photographed after 24 hours. The relative survival of each of these strains was calculated with the following formula. *Relative viability = (CFU-1 ml of treatment/ CFU-1 ml of control).* Results obtained from at least three independent sets of experiments are represented as graphs. Appropriate antibiotic(s) were added in the growth medium when plasmid-containing strains were used.

### Recombinant DNA techniques

Procedures for DNA isolation, amplification, PCR purification, agarose gel electrophoresis, and transformation of *E. coli* were performed as described by (50). Genomic DNA and plasmid DNA were isolated and purified using the Wizard® Genomic DNA Purification Kit and PureYield™ Plasmid Miniprep System (Promega, USA) according to the manufacturer’s instructions. Wizard® SV Gel and PCR Clean-Up System (Promega, USA) was used for PCR and gel purification according to the manufacturer’s instructions. Enzymes were purchased from New England Biolabs and used as described by the manufacturer. For PCR amplification, Phusion® High Fidelity PCR Kit (NEB, England) were used.

### Microscopy

Live cell imaging was done as described previously (26). 1 ml of bacterial culture was centrifuged and dissolved in 0.5ml of phosphate-buffered saline and was spotted on 1% agarose pads (in 0.85% saline) with uncoated coverslips before visualization under a microscope. To check the complementation effect of periplasmic proteases, log-phase cells were treated with A22 (2µg/ml), with subsequent induction using 0.01% arabinose. Cells were visualized and photographed using a Nikon Eclipse Ti2-E equipped with a 100X CFI Plan Apochromat oil objective and a DSQi-2 Monochrome Camera (Nikon). Images were processed using the NIS Elements AR software (Nikon).

## Declarations

### Author contribution

Conceptualization, S.G, O.A-C; Methodology, T.N, M.N, B.K, S.K; Data Analysis, M.N, S.M; Investigation, T.N, M.N; Writing – Original Draft, T.N, M.N, S.G; Writing – Review & Editing, S.G, O.A-C.; Resources, S.G, O.A-C.; Funding Acquisition, S.G., O.A-C.

### Availability of data and material

All supporting data are included in the additional files. The materials generated as part of this study are available upon request.

## Acknowledgment

Sutharsan Govindarajan acknowledges support from DST-SERB (CRG/2020/003295) and the DBT-Wellcome Trust Early Career Fellowship (IA/E/19/1/504958). Research in the Amster-Choder lab was supported by the Israel Science Foundation (ISF) founded by the Israel Academy of Sciences and Humanities (grant no. 1274/19). OAC is an incumbent of the Dr. Jacob Grunbaum Chair in Medical Sciences. T. Nagarajan acknowledges a Postdoctoral Fellowship from SRM University - AP. All authors appreciate the infrastructure support received from SRM University – AP, India (SRMAP/URG/GENERAL/2023-24/010 and SRMAP/URG/E&PP/2022-23/018). We are grateful for helpful discussions with members of the Department of Biological Sciences, SRM University – AP.

## SUPPLEMENTARY FIGURE AND TABLE LEGENDS

**Table S1 –** Bacterial Strains and Plasmids used in this study

**Table S2 –** Primers used in this study

**Table S3 –** Transcriptome analysis Raw data

**Table S4 –** Differentially expressed genes in *ΔmreBCD* compared to WT, A22 compared to untreated, and those commonly regulated in both conditions

**Table S5 –** Summary of results of KEGG Pathway enrichment analysis

**Fig S1 –** Relative viability of wild-type E. coli cells grown in the presence of different concentrations of A22

**Fig S2 –** Serial dilutions of indicated strains on LB plates containing 0.05% arabinose and grown at 30°C or 42°C overnight.

