## Supplementary Figures for "Periplasmic proteostasis enables bacterial survival during MreB cytoskeletal disruption"

**Fig S1.**

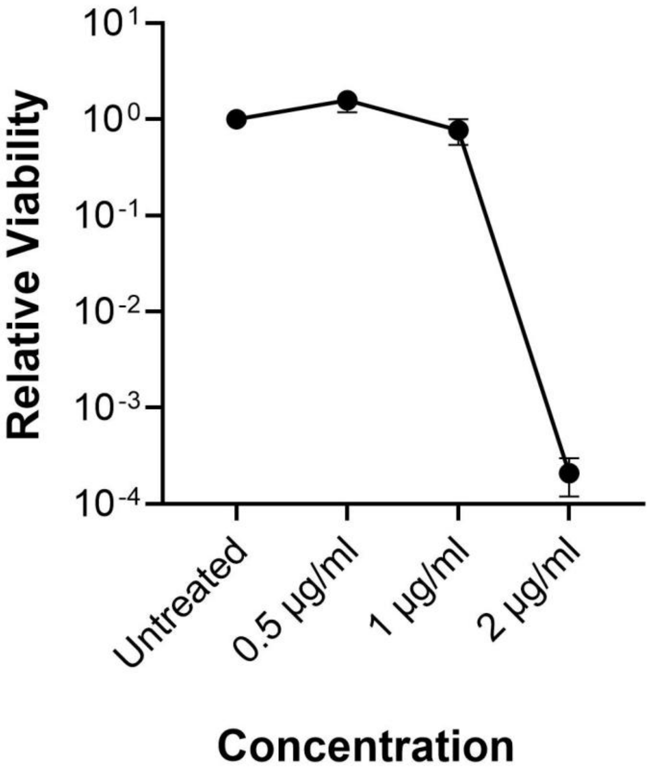


**Fig S1 – Relative viability of wild-type *E. coli* cells grown in the presence of different concentrations of A22.** Early-log-phase cultures of wild-type cells were spotted onto plates containing various concentrations of A22 or onto untreated control plates and incubated overnight. Relative viability was calculated as described in the Methods. The graph shows the mean of three independent experiments, with error bars indicating variability.

### Fig S2.

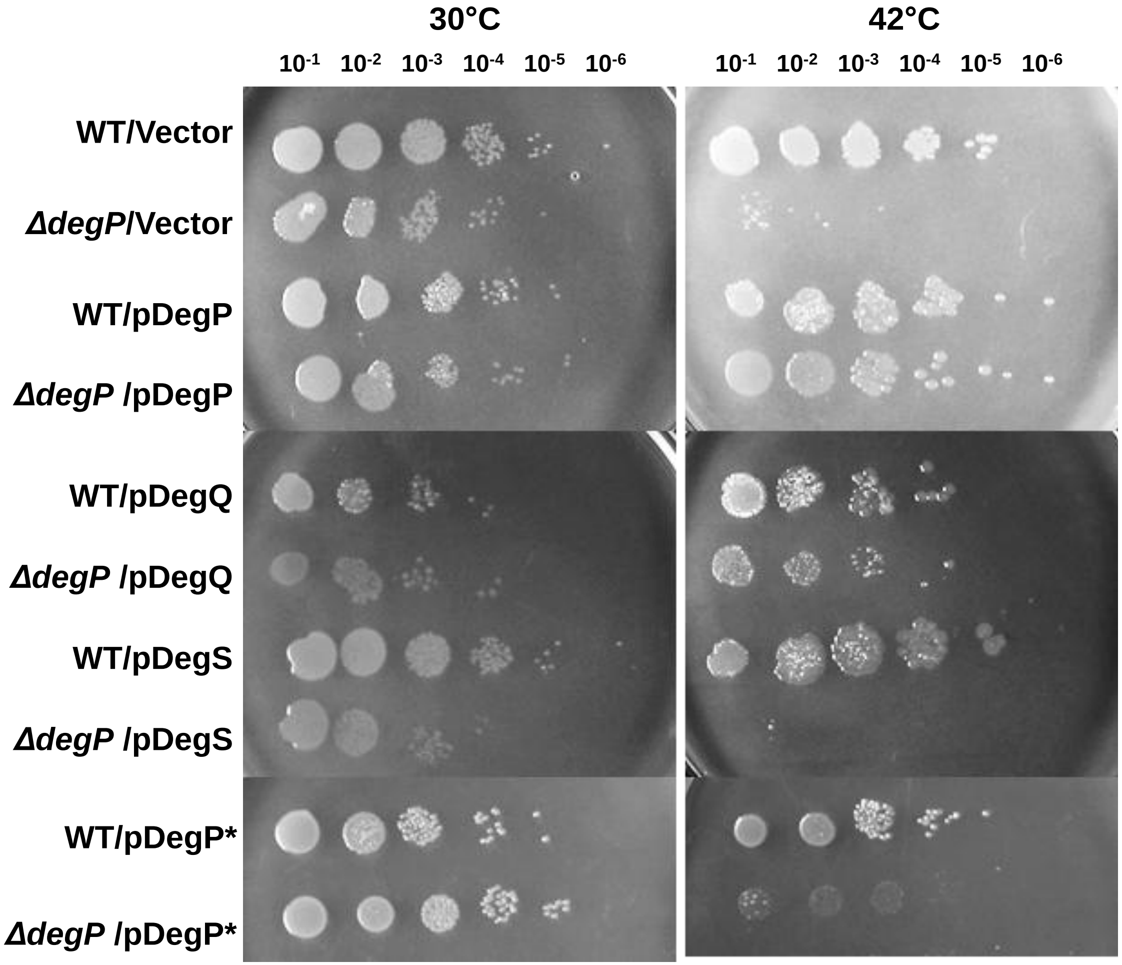


**Fig S2 – Serial dilutions of the indicated strains on LB plates containing 0.05% arabinose and grown at 30°C or 42°C overnight.**
